# Assessment of mosquito longevity, fecundity, and dengue virus titer following exposure to plasma from a host with Type 2 diabetes mellitus – a pilot study

**DOI:** 10.1101/2022.07.28.501853

**Authors:** Chelsea T. Smartt, Natalie Kendziorski, Tse-Yu Chen, Sara Farless

## Abstract

The mosquito *Aedes aegypti* (Linnaeus) prefers feeding on humans and will encounter many blood components that might influence mosquito physiology and vectorial capacity. The effects of dengue virus infection in patients with type-2 diabetes mellitus revealed an influence on platelet numbers in blood and diabetic persons are more likely to develop severe dengue. Infection with dengue virus (DENV) is known to impact diabetic patients but the impact of diabetic blood on mosquito vectorial capacity is not known. This pilot study investigated the effects of ingesting blood plasma from a patient with diabetes on mosquito fecundity, longevity, and vector competence for DENV-2 in the mosquito, *Ae. aegypti*. Female mosquitoes fed on plasma from a diabetic individual laid significantly more eggs. Diabetic plasma shortened overall mosquito life span. Vector competence was significantly different between extrinsic incubation periods at 7 (3.44±2.5 log PFUe DENV) and 14 days post-infection (4.28±2.4 log PFUe DENV) in mosquitoes fed DENV-2 in plasma from a diabetic individual. As this study is an initial investigation into the influence of diabetes plasma on mosquito life history traits, a more comprehensive study with more samples of diabetes plasma is underway and should yield more details about the impact of diabetes on mosquito biology.

## Introduction

Mosquito-borne diseases are major health concerns worldwide. This is due in part to emerging and re-emerging mosquito borne diseases caused by viruses, most recently dengue, Zika and chikungunya viruses [Mayer et al. 2017]. *Aedes aegypti* (L.) prefers to feed on human blood and has enabled this species to spread many human disease-causing pathogens [Crawford et al. 2017, Rose et al. 2020].

Acquisition of blood from humans means encountering many blood-borne molecules that influence mosquito physiology and mosquito vector competence. Blood from a human with blood diseases or taking certain medications may impact mosquito biology [Gendrin et al. 2015]. The number of people with abnormal levels of glucose in their blood, as with diabetes disease, has increased worldwide [Saeedi et al. 2019]. Diabetes is one of the top 10 diseases that kill adults worldwide, and whose prevalence is expected to go from 9.3% in 2019 to 10.3% by 2030 [Wild et al. 2004]. With the predicted increase in diabetic people worldwide, the impact on mosquito biology, especially for *Ae. aegypti*, should be determined [Harrington et al. 2001, Powell 2018]. High glucose e in the blood may alter mosquito biology and ability to transmit viruses [Weng et al. 2021]. Investigation of these effects will provide information about possible mosquito biology adaptations.

Patients with Type-2 diabetes mellitus may develop severe dengue [Chen et al. 2015]. However, Latt et al. [2020] report that although diabetes patients were more likely to get severe dengue, there was no support for diabetes being an indicator of severity of DENV disease. This pilot study investigated the effects of ingesting blood plasma from a patient with diabetes on mosquito longevity, fecundity, and vector competence for DENV-2 in *Ae. aegypti*.

## Materials and Methods

A Vero Beach, FL population of *Ae. aegypti* mosquitoes (F3) were held at 28°C and 60-80% relative humidity with a light: dark cycle of 14:10 hours. The eggs were hatched under vacuum and reared following established methods [Chen et al. 2021a]. Adult female mosquitoes were mated prior to experimentation and sugar-starved 24 hours before being offered blood or infectious blood.

Dengue-2 virus strain New Guinea C (DENV-2, GenBank Accession # KM204118) was propagated in African green monkey (Vero) cells in M199 medium (Gibco, Waltham, MA, USA) at a multiplicity of infection of 0.01 viruses per cell following established protocols [Chen et al. 2021a].

Fifty female *Ae. aegypti* (4-6 days old) were transferred into cardboard cartons, covered with mesh (Solo Cup Company, Illinois, US). Four replicate cartons were used per treatment (200 females/ treatment). For the infectious blood-feeding experiment, defibrinated bovine blood (Hemostat Laboratories, Dixon, CA) was warmed at 37°C in a water bath. Commercially available plasma from non-diabetic people (control, pooled from 8 individuals) and plasma from a diabetic female was purchased (ZenBio Inc. Durham, NC, US). Plasma from blood can be absorbed by *Ae. aegypti* for egg production and therefore assessment of change in life history traits due to diabetes disease is possible using commercially available plasma [Gonzales et al. 2015]. A single source of diabetes plasma was available for purchase from ZenBio and all plasma samples are screened for infectious pathogens (www.zen-bio.com).

Blood meals contained 1 ml bovine blood with 100mM adenosine triphosphate, 1 ml of DENV-2 and either 1 ml human regular plasma or 1 ml of human Type 2 diabetes plasma. *Ae. aegypti* were fed blood meals using the Hemotek membrane feeding system (Hemotek, Blackburn, UK) at 28°C in an incubator in a Biosafety Level 2 facility. Engorged females were sorted, counted and returned to the cardboard cartons by treatment group. Engorged mosquitoes were provided 5% sucrose solution *ad libitum* and maintained in an incubator as described (Chen et al. 2021a). Daily mortality rates post-infection were checked and dissection and salivation using capillary tubes was completed at 7- and 14-days post-infection (dpi) to determine dengue virus dissemination and transmission as described [Anderson et al. 2010]. Bodies, legs and saliva samples collected to evaluate virus titer were stored in 1.5 ml Eppendorf tubes at −80°C until processing.

The same cohort of *Ae. aegypti* was used for life history trait estimation. The longevity of mosquitoes that fed on the same batches of commercially available human plasma was assessed. Moistened brown seed germination paper was inserted into each vial (100 vials per treatment) to serve as an oviposition surface. A gravid female was introduced into each vial and the opening secured with a sugar-dampened cotton plug. Cotton was rehydrated as needed and vials maintained in similar conditions as described. Survival rates post-feeding were recorded on day one, three, and seven, although observations of survivorship and fecundity continued every day for one month until the final mosquito died. Total viable eggs were counted under a Leica stereo microscope (TechniQuip Corp., Pleasonton, CA, US). The Mann-Whitney test was used to calculate the p-value in the fecundity study and Log-rank (Mantel-Cox) test was applied in the longevity experiment.

Mosquitoes were collected 7 days post-blood feeding from control and diabetes treatment groups (four replicates each) and five female mosquitoes were combined in each tube. TRIzol Reagent (Invitrogen, Carlsbad, CA, USA) was used to extract RNA from mosquito samples following well-established protocols [Smartt et al. 2017]. The transcript levels were determined by Bio-Rad CFX96™ Real-Time PCR using iTaq™ Universal SYBR Green One-Step Kit (Bio-Rad, Hercules, CA, USA) and primers designed to select innate immune response pathways genes (Table S1). *Ae. aegypti* ribosomal protein S7 gene (GenBank Accession # AY380336) was used as a control for standardizing transcript levels. The gene expression in each group was compared by delta-delta Ct value analysis between control and diabetes experiment groups. Wilcoxon test was used to calculate p-values and only p-value < 0.05 was counted as significantly different.

To determine the infection, dissemination, transmission rates and virus genome equivalents in the treatment group mosquitoes, mosquito body, leg, and saliva RNA from individual female mosquitoes was used in qRT-PCR reactions with DENV-2 specific primers, iTaq Universal SYBR Green One-step kit and analyzed on a Bio-Rad PCR system as described. Viral genome equivalents were calculated based on a standard curve and recorded as log (base 10) plaque-forming unit equivalents per milliliter (log PFUe/ml). The standard curves for DENV genome equivalents were described previously [Chen et al. 2021a].

## Results

### Fecundity

Mated female *Ae. aegypti* were exposed to commercially purchased human plasma, from individuals with or without Type 2 diabetes, and eggs laid per individual female counted. Female mosquitoes that ingested plasma from a diabetic individual laid significantly more eggs (40.44±4.74 eggs/female, 4075 eggs total, p-value=0.02) than those exposed to plasma from non-diabetic people (31.26±3.79 eggs/ female, 3192 eggs total, Figure 1) although the feeding rates were the same (66.67%). Fewer mosquitoes fed on control plasma (19 out of 100 females) laid more than 50 eggs compared to individual females fed plasma from a diabetic individual (43) although the proportion of egg-laying females per treatment was higher for mosquitoes fed control plasma (96.77±1.15%) compared to those fed diabetic plasma (91.06±6.14%).

**Figure 1.**
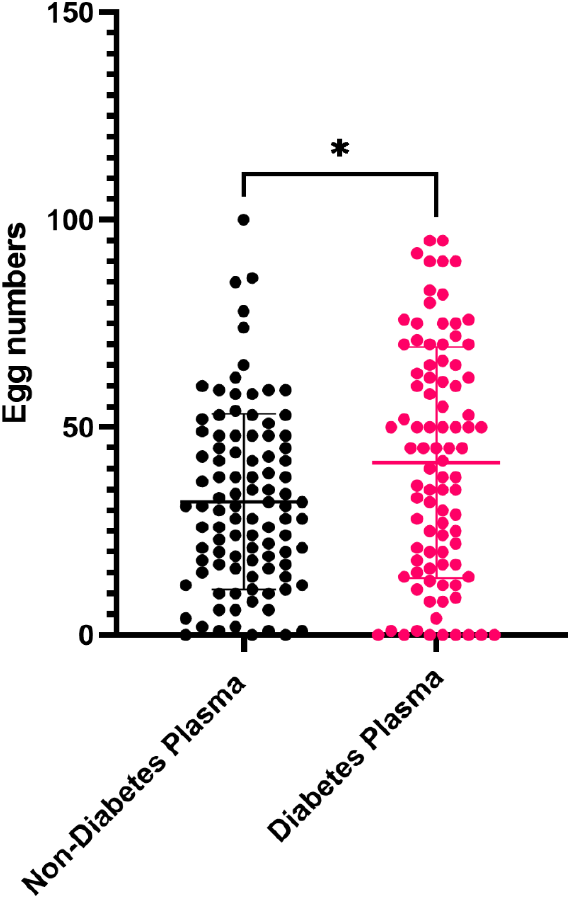
Fecundity is significantly affected in *Ae. aegypti* fed plasma from a person with diabetes (Diabetes) and non-diabetic individuals (Control) (Mann-Whitney test, *P = 0.02).

### Longevity

Female mosquitoes fed plasma from a diabetic person showed delayed mortality with little death occurring before day 18, and ∼50% of the death occurring at day 19 (Figure 2). Mortality for female mosquitoes fed control plasma started at day 10 and reached ∼50% by day 18. The mosquitoes fed control plasma lived longer (36 days) compared to mosquitoes fed plasma from a diabetic person (32 days), however the difference was not significant (Log-rank test: p-value=0.1748; Gehan-Breslow-Wilcoxon test: p-value=0.1578).

**Figure 2.**
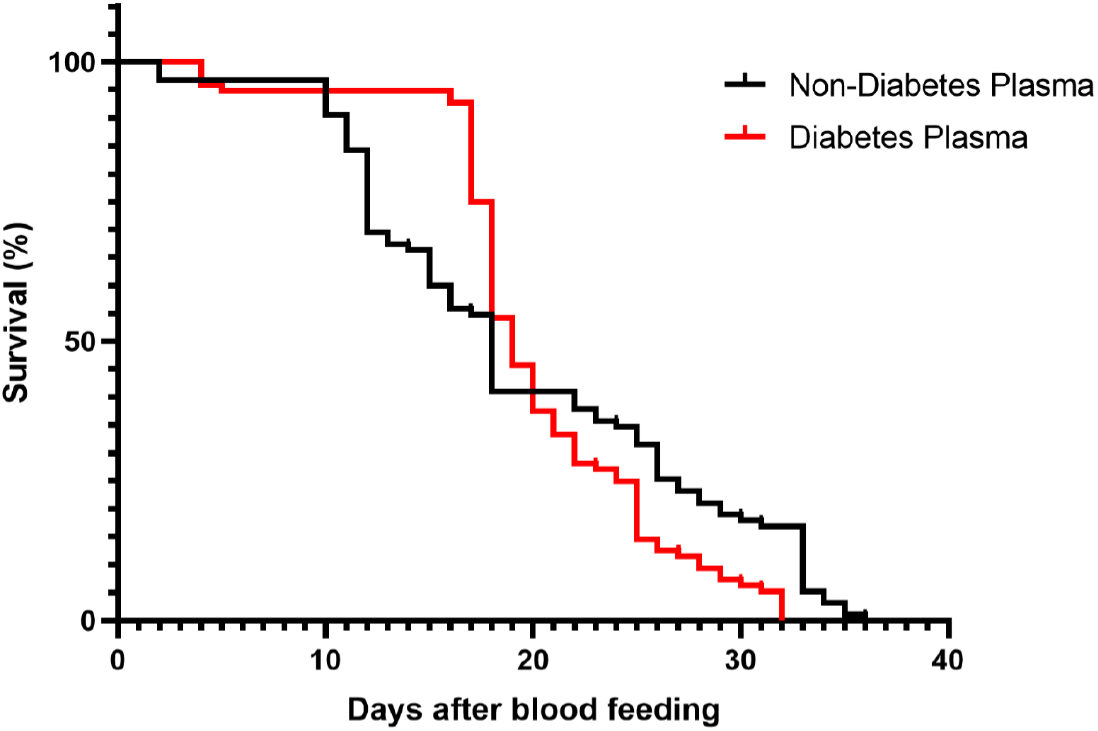
Longevity is shorter in female *Ae. aegypti* fed plasma from a person with diabetes (Diabetes) compared with non-diabetic individuals (Control).

### Vector competence for DENV-2

The effects of commercially purchased diabetic plasma on DENV-2 vector competence was investigated in the same *Ae. aegypti* population by exposing mosquitoes to 8.44± 0.44log PFUe DENV-2. The mosquitoes ingested similar titers of DENV-2 (mosquitoes fed diabetic plasma = 5.84±0.66 log PFUe; fed control plasma = 5.28±0.73 log PFUe, p-value=0.7). At 7dpi, the infection, dissemination and transmission rates were not significantly different (Figure 3A). The titers of DENV-2 in mosquito bodies, legs and saliva between those fed diabetic plasma compared to mosquitoes fed control plasma were not significantly different. Also transmission rates were low at 7dpi with only a few mosquitoes fed the control plasma showing virus in their saliva (11%, Figure 3A). At 14 dpi, dissemination and transmission rates did not significantly differ nor did the DENV-2 titers differ between the treatment groups. More mosquitoes fed control plasma were positive for DENV-2 in saliva (Figure 3B). Within population comparison of titers between extrinsic incubation periods (7 and 14 dpi) showed mosquitoes fed DENV-2 in plasma from a diabetic individual were significantly different between 7 (3.44±2.5 log PFUe DENV) and 14 dpi (4.28±2.4 log PFUe DENV, p-value=0.046), compared to mosquitoes fed DENV-2 in control plasma (Figure 3A and B).

**Figure 3.**
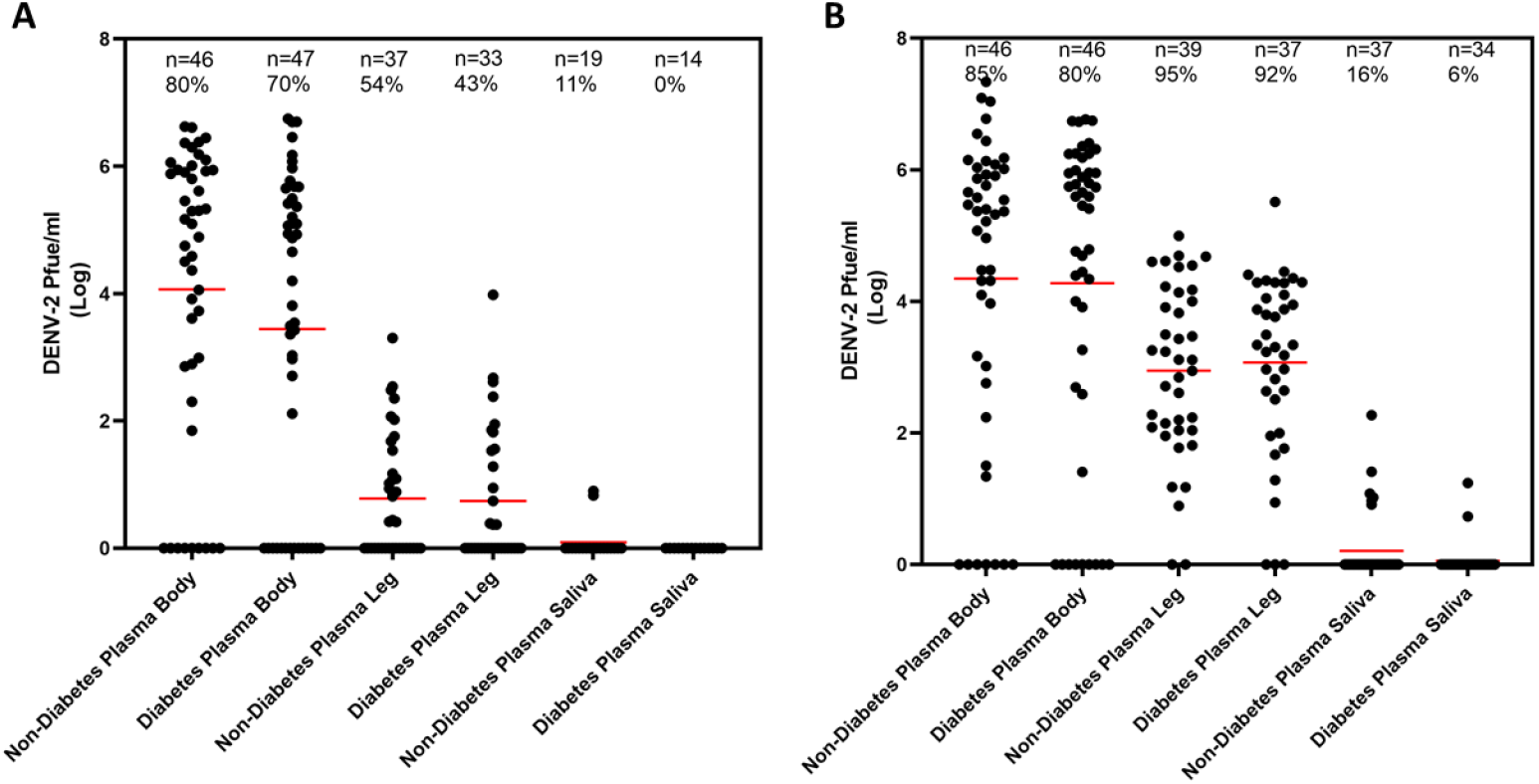
Vector competence for dengue virus serotype 2 (DENV-2) at (A) 7 days (B) 14 days post infection in bodies, legs and saliva collected from adult female *Ae. aegypti* fed plasma from a per-son with diabetes (Diabetes) compared with plasma from non-diabetic individuals (Non-Diabetes). DENV = dengue virus, dpi = days post infection, % = infection, dissemination and transmission rates, n = number of mosquitoes analyzed.

### Expression of innate immune response genes

The role of the innate immune response in lower DENV-2 titer in bodies at 7dpi in mosquitoes fed plasma from a diabetic person was assessed. Expression of dome (Janus kinase/signal transducers and activators of transcription), Rel-1A and Rel-1B (Toll pathway) and Rel-2 (immune deficiency pathway) was detected in DENV-infected mosquitoes collected at 7dpi after ingestion of both plasma from a diabetic individual and control plasma, however there were no expression differences (Figure 4, Table S2). Expression of the Ago-2 gene, involved in the RNAi pathway, was significantly lower in mosquitoes fed plasma from a diabetic individual (fold change = 0.52±0.14) compared to those who ingested control plasma (fold change = 1.01±0.14, p-value =0.0286, Figure 4).

**Figure 4.**
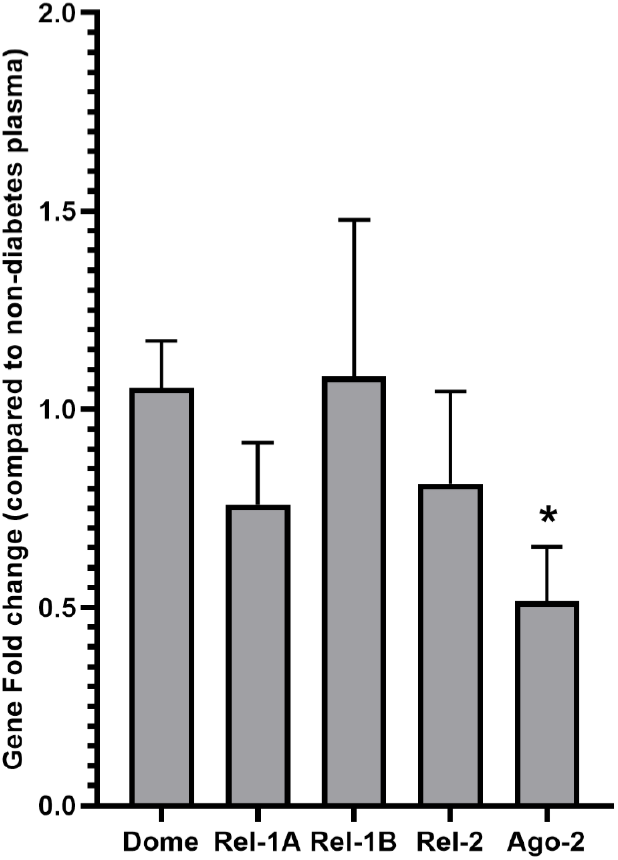
Expression changes in select immune response genes in adult female *Ae. aegypti* fed plasma from a person with diabetes containing DENV-2 compared with plasma from non-diabetic individuals containing DENV-2. (Wilcoxon, *P =0.0286).

## Discussion

Mosquito borne diseases pose a substantial human health concern globally, as it disrupts everyday human health and impacts overall quality of life [Franklinos et al. 2019]. With a decrease in effective mosquito control methods, wild mosquitoes will encounter humans more frequently for blood feeding [Chen et al. 2021b, McGregor and Connelly 2021], including individuals with type 2 diabetes mellitus known to be more susceptible to vector borne diseases [Sorenson et al. 2016, Lee et al. 2020]. Considering the importance of understanding female mosquito physiology on controlling mosquito borne viruses, longevity, fecundity, and vector competence for DENV-2 in mosquitoes fed diabetes plasma were assessed and preliminary findings reported.

The number of female *Ae. aegypti* that fed on plasma from a human with diabetes did not differ from the number that fed on control plasma. Female *Ae. aegypti* fed plasma from a diabetic person laid significantly more eggs than those fed control plasma. This suggests that blood containing sugar might support higher egg production [Wu et al. 2021]. A previous study showed that *Ae. aegypti* fed readily on plasma from humans and plasma supported development of eggs [Harrison et al. 2021]. Therefore, the differences in fecundity between female *Ae. aegypti* in the treatment groups is interesting but could be due to high levels of glucose. Estimates of longevity for female *Ae. aegypti* fed plasma showed no significant differences between treatments, which was also consistent with what has been reported in the literature [Scott et al. 1997]. The effects of feeding on plasma from a diabetic individual on mosquito vector competence for DENV-2 revealed modest variation between treatment groups. Mosquitoes fed control plasma had higher infection rates at 7dpi and the DENV titers were higher than in mosquitoes that were fed on diabetic plasma. Lower DENV-2 titers in mosquitoes fed plasma from a diabetic person might be attributed to glucose-induced bacterial growth and blood iron content [Wang et al. 2011, Zhu et al. 2019]. The expression of established immune response pathway genes that might have a role in controlling DENV titers at 7 dpi showed no significant alterations between the treatment groups [Carvalho-Leandro et al. 2012]. The expression of the Ago-2 gene was significantly lower in mosquitoes fed diabetes plasma containing DENV-2 compared to the control group. The reduced expression of Ago is surprising as sugar feeding was shown to not affect Ago-2 expression. Perhaps the decrease in Ago-2 expression is due to presence of gut bacteria, as expression of the Ago-2 differed between non-sugar fed and sugar fed aseptic mosquitoes [Almire et al. 2021]. This study revealed that that some life history traits in *Aedes aegypti* may alter when feeding on a patient with diabetes.

## Acknowledgments

We thank Dr. Jasbir Singh for intellectual discussions. The authors would like to thank FMEL and NSF CAMTech for support.

## Institutional Review Board Statement

Not applicable.

## Author Contributions

Conceptualization, C.T.S.; methodology, N.K., S.O.; validation, N.K., S.O.; and T-Y.C.; formal analysis, T-Y.C.; investigation, N.K., S.O.; resources, C.T.S.; data curation, C.T.S. and T-Y.C.; writing—original draft preparation, C.T.S.; writing—review and editing, N.K., T-Y.C.; visualization, T-Y.C.; supervision, C.T.S.; project administration, C.T.S.; funding acquisition, C.T.S. All authors have read and agreed to the published version of the manuscript.

## Supplementary Materials

Table S1: Primer sequences for gene expression and detection of virus;

Table S2: Relative gene expression of select immune response genes.

