## Supplementary material for "Assessment of mosquito longevity, fecundity, and dengue virus titer following exposure to plasma from a host with Type 2 diabetes mellitus – a pilot study": Table S1

| **Name** | **Sequence** |
| --- | --- |
| F- Dome | 5’- GCA TCA GCG GGA AAG TTC CAA TGT -3’ |
| R- Dome | 5’-AGC TTG TAA TCG GTG GGA ATC GGZ-3’ |
| F- Argonaute-2 | 5’-ACA ACA GCA ACA ATC CCA GA-3’ |
| R- Argonaute-2 | 5’-GTG GAC GTT GAT CTT GTT GG-3’ |
| F- Rel-1A | 5’-AGA AAAG CCA TGT CCG ATC TGGTGA-3’ |
| R- Rel-1A | 5’-CCT GTT TGT GCA CGT TGGTAT GCT-3’ |
| F- Rel-1B | 5’-AAA CTT CCT CTG CCT CCC AAA-3’ |
| R- Rel-1B | 5’-TAC GCA TGG AAC CCT TCC GAA TGA-3’ |
| F- Rel-2 | 5’-GGACTGGGGTTCTTTCTCGG-3’ |
| R- Rel-2 | 5’-ATTTGTCTCGTGGCCGGTAG-3’ |
| DENV-2 F | CTWTCAATATGCTGAAACGCG |
| DENV-2 R | CGCCACACAAGGGCCATGAACAG |
| F- S7 | 5’- ACAAGAACCAGCAGACCAC -3’ |
| R- S7 | 5’- TCCGGGAATTCGAACGTAAC -3’ |

Table S1: Primer sequences for gene expression and detection of virus
