## Supplementary material for "Assessment of mosquito longevity, fecundity, and dengue virus titer following exposure to plasma from a host with Type 2 diabetes mellitus – a pilot study": Table S2

Table S2: Relative gene expression of select immune response genes in mosquito bodies infected with DENV in plasma from non-diabetic individuals and DENV in plasma from a diabetic person at 7dpi.

| Gene | Control | Diabetes |
| --- | --- | --- |
| Dome | 1.01±0.14 | 1.06±0.12 |
| Rel-1A | 1.10±0.46 | 0.76±0.16 |
| Rel-1B | 1.02±0.22 | 1.08±0.40 |
| Rel-2 | 1.02±0.23 | 0.81±0.23 |
| Ago-2 | 1.01±0.14 | 0.52±0.14 |
